# Branched germline cysts and female-specific cyst fragmentation facilitate oocyte determination in mice

**DOI:** 10.1101/2022.11.17.516957

**Authors:** Kanako Ikami, Suzanne Shoffner, Malgorzata Gosia Tyczynska Weh, Santiago Schnell, Jingqun Ma, Shosei Yoshida, Edgar Diaz Miranda, Sooah Ko, Lei Lei

**Affiliations:** Department of Cell and Developmental Biology, University of Michigan Medical School, Ann Arbor, Michigan, 48109; Buck Institute for Research on Aging, Novato, California, 94945; Department of Molecular & Integrative Physiology, University of Michigan Medical School, Ann Arbor, Michigan, 48109; The Bioinformatics Core, University of Michigan Medical School, Ann Arbor, Michigan, 48109; Division of Germ Cell Biology, National Institute for Basic Biology, Okazaki 444-8585 Aichi, Japan; Department of Basic Biology, School of Life Science, SOKENDAI (Graduate University for Advanced Studies), Hayama, Kanagawa 240-0193, Japan; Department of Obstetrics, Gynecology and Women’s Health, University of Missouri School of Medicine, Columbia, Missouri, 65211; Division of Biological Sciences, College of Arts and Sciences, University of Missouri, Columbia, Missouri, 65211; Department of Microbiology & Molecular Genetics, University of California, Davis, Davis, California, 95616; Department of Biological Sciences and Department of Applied and Computational Mathematics and Statistics, University of Notre Dame, Notre Dame, Indiana, 46556; Pathology Department, St. Jude Children’s Research Hospital, Memphis, Tennessee, 38105

## Abstract

During mouse gametogenesis, germ cells derived from the same progenitor are connected via intercellular bridges forming germline cysts, within which asymmetrical or symmetrical cell fate occurs in female and male germ cells respectively. Here, we have identified branched cyst structures in mice, and investigated their formation and function in oocyte determination. In fetal female cysts, 16.8% of the germ cells are connected by three or four bridges, namely branching germ cells. These germ cells are preferentially protected from cell death and cyst fragmentation, and to accumulate organelles and cytoplasm from sister germ cells to become primary oocytes. Changes in cyst structure and single-cell mRNA profiles suggested that cytoplasmic transport in germline cysts is conducted in a directional manner, in which cellular content is first transported locally between peripheral germ cells and further enriched in branching germ cells, a process causing selective germ cell loss in cysts. Cyst fragmentation occurs extensively in female cysts, but not in male cysts. Male cysts in fetal and adult testes have branched cyst structures, without differential cell fates between germ cells. During cyst formation, E-cadherin junctions between germ cells position intercellular bridges to form branched cysts. Disrupted junction formation in E-cadherin-depleted cysts led to an altered ratio in branched cysts. Germ cell-specific E-cadherin knockout resulted in reductions in primary oocyte number and oocyte size. These findings shed light on the mechanism of how the size of the ovarian reserve, the number of primary oocytes available to sustain adult ovarian function, is determined during ovary formation.

## Introduction

In mammals, germline differentiation during gametogenesis produces eggs and spermatozoa that are distinct in morphology and function, yet are derived from the same type of progenitors, primordial germ cells (PGCs) during fetal gonad development (1). In the fetal testis, PGCs differentiate into gonocytes, the majority of which differentiate into spermatogonia stem cells (SSCs) to sustain spermatogenesis in the adult testis (2–4). In the fetal ovary, PGCs differentiate into primary oocytes, which become quiescent along with the surrounding pregranulosa cells, forming primordial follicles. The pool of dormant primordial follicles serves as the ovarian reserve that sustains egg production and ovarian function in adulthood (5, 6). An unresolved question in mammalian oocyte differentiation is the differential cell fate among germ cells during this process. In both humans and mice, only a small fraction (~15-20%) of fetal germ cells survive to become primary oocytes, with the majority of the germ cells undergoing cell death (7, 8). Mechanisms underlying the distinct cell fates during oocyte differentiation are key to understanding ovarian reserve formation and associated ovarian health issues.

During gametogenesis in both female and male gonads, germ cells form a highly conserved cellular structure, the germline cyst. Within the cyst, sister germ cells derived from a single progenitor are connected via intercellular bridges formed through incomplete cytokinesis during mitotic divisions. Germline cysts are found in invertebrates, such as *Drosophila*, and vertebrates, including mice and humans (9–14). During mouse gametogenesis, PGCs form germline cysts in the fetal ovary and testis from embryonic day 10.5 (E10.5) to E14.5. Female germ cells enter meiosis immediately following mitotic cyst formation, while male germ cells undergo G0/G1 arrest at E14.5 (1, 11). Intercellular bridges have been observed by electron microscopy and can be detected by antibody staining of specific markers, such as TEX14 (testis-expressed 14) and RacGAP (Rac GTPase Activating Protein) in fetal ovaries, as well as fetal and adult testes (8, 14–16). In adult testes, as SSCs initiate spermatogenesis, male germ cells undergo mitosis to form germline cysts. Interconnected spermatocytes enter meiosis synchronously as they differentiate into spermatozoa (15, 17).

A conserved phenomenon of gametogenesis is asymmetric cell fate in female germline cysts. During oogenesis in *Drosophila* and mice, organelle and cytoplasmic transport between sister germ cells within a cyst facilitates two distinct cell fates: becoming oocytes vs. undergoing cell death (18, 19). Within each *Drosophila* 16-cell cyst, one of the two germ cells that are connected by four intercellular bridges progresses through meiosis and differentiates into an oocyte by collecting organelles and cytoplasmic content from the remaining 15 sister germ cells, i.e. nurse cells. The differential cell fate in the *Drosophila* cyst is facilitated by the fusome, a cytoplasmic organelle that asymmetrically spans through the intercellular bridges during cyst formation, with the future oocyte retaining a greater proportion compared to the nurse cells (19–21). The fusome anchors germ cell mitotic spindles causing branched cyst formation and organizes microtubule minus ends into the future oocyte, facilitating directional organelle and cytoplasmic transport from the nurse cells to the future oocyte (19–22).

During mouse oocyte differentiation that takes place from E14.5 to postnatal day 4 (P4), germline cysts fragment gradually and become individual cells. On average, 20% of the E14.5 cyst germ cells differentiate into primary oocytes through organelle and cytoplasmic collection within germline cysts. The remaining 80% of the E14.5 cyst germ cells undergo cell death. During this process, an average four-fold increase in organelle content and cell volume occurs in primary oocytes (11, 18). In the primary oocytes, centrosomes, Golgi complexes, endoplasmic reticulum and mitochondria organize into a Balbiani body (B-body), in which the Golgi complexes have a characteristic spherical arrangement (18, 23). The fusome structure is not observed in mouse cysts by EM (8). How germ cells are connected structurally within the mouse cyst and how specific germ cells are selected to receive organelles and cytoplasmic content to become primary oocytes remain open questions.

During spermatogenesis in *Drosophila* and adult mice, intercellular bridges facilitate the transfer of signaling molecules between cyst germ cells, enabling synchronized meiosis and symmetrical cell fate in the cyst (15, 24). All male germ cells in the germline cyst progress through spermatogenesis to differentiate into sperm or otherwise undergo cell death when triggered in one germ cell (25–28). The *Drosophila* male cyst is in the same branched structure as the female cyst, with the fusome evenly distributed in each cyst germ cell (29, 30). The structures of male germline cysts in fetal and adult mouse testes have yet to be characterized.

In the present study, we find that both female and male mouse germline cysts are in branched structures. On average, 16.8% of the germ cells in E14.5 ovaries are connected by three or four intercellular bridges, namely branching germ cells. These germ cells and the associated bridges are preferentially protected from cell death and cyst fragmentation, receiving organelles and cytoplasm to become primary oocytes. Male cysts in fetal and adult testes have branched cyst structures without apparent differential cell fates. We further demonstrated that during cyst formation from single PGCs, E-cadherin junctions between sister germ cells determine the position of intercellular bridges and lead to the formation of branched cysts, and the loss of E-cadherin in germ cells results in altered ratio in branched cysts and defective oocyte differentiation. In summary, our present study demonstrates that branched cyst structure is involved in determining oocyte fate in the germline cyst during oocyte differentiation. These findings shed light on the mechanism of how the size of the ovarian reserve, the number of oocytes available to sustain adult ovarian function, is determined during ovary formation at the fetal stage.

## Results

### Terms and definitions

#### Intercellular/connected bridge

the TEX14 (or RacGAP)-positive ring or puncta at the size of approximately 1 μm connecting two germ cells.

#### Unconnected bridge

the TEX14 (or RacGAP)-positive ring or puncta at the size of approximately 1 μm that is connected to one germ cell. The unconnected bridge may represent the remains of an intercellular bridge.

#### Germ cell clone

germ cells derived from a single lineage-labeled primordial germ cell, regardless of whether they remain connected by intercellular bridges.

#### Germline cyst

a cluster of interconnected germ cells derived from the same progenitor germ cell and generated by mitotic divisions with incomplete cytokinesis.

#### Branching germ cell

the germ cell that is connected by three or four intercellular (or unconnected) bridges in the germline cyst.

#### Branched germline cyst

the germline cyst that contains one or more branching germ cells.

### Female and male PGCs form similar structured branched germline cysts

To elucidate the structure of mouse cysts, i.e. geometrical locations of germ cells and bridges in the cyst, during cyst formation, as well as the subsequent changes during gametogenesis in fetal gonads, we conducted single-cell lineage tracing to label individual PGCs at E10.5 by injecting a single low dose of tamoxifen into pregnant female *R26R-YFP* mice mated with male *CAG-creER; R26R-YFP* mice (11) (Fig. 1A). This approach allows us to follow the differentiation of the germ cell clones derived from individual lineage labelled PGCs, which are comprised of various numbers of germline cysts, by the expression of lineage marker YFP (Fig. 1B). Fetal ovaries were collected at E12.5, representing the cysts in the middle of cyst formation; and E14.5, E17.5 and P0, representing the cysts prior to, during, and at the end of organelle transport, respectively. Fetal testes at these four time points were also collected for a comparative analysis on cyst structure during gonocyte differentiation (Supplementary Table 1-8).

**Figure 1.**
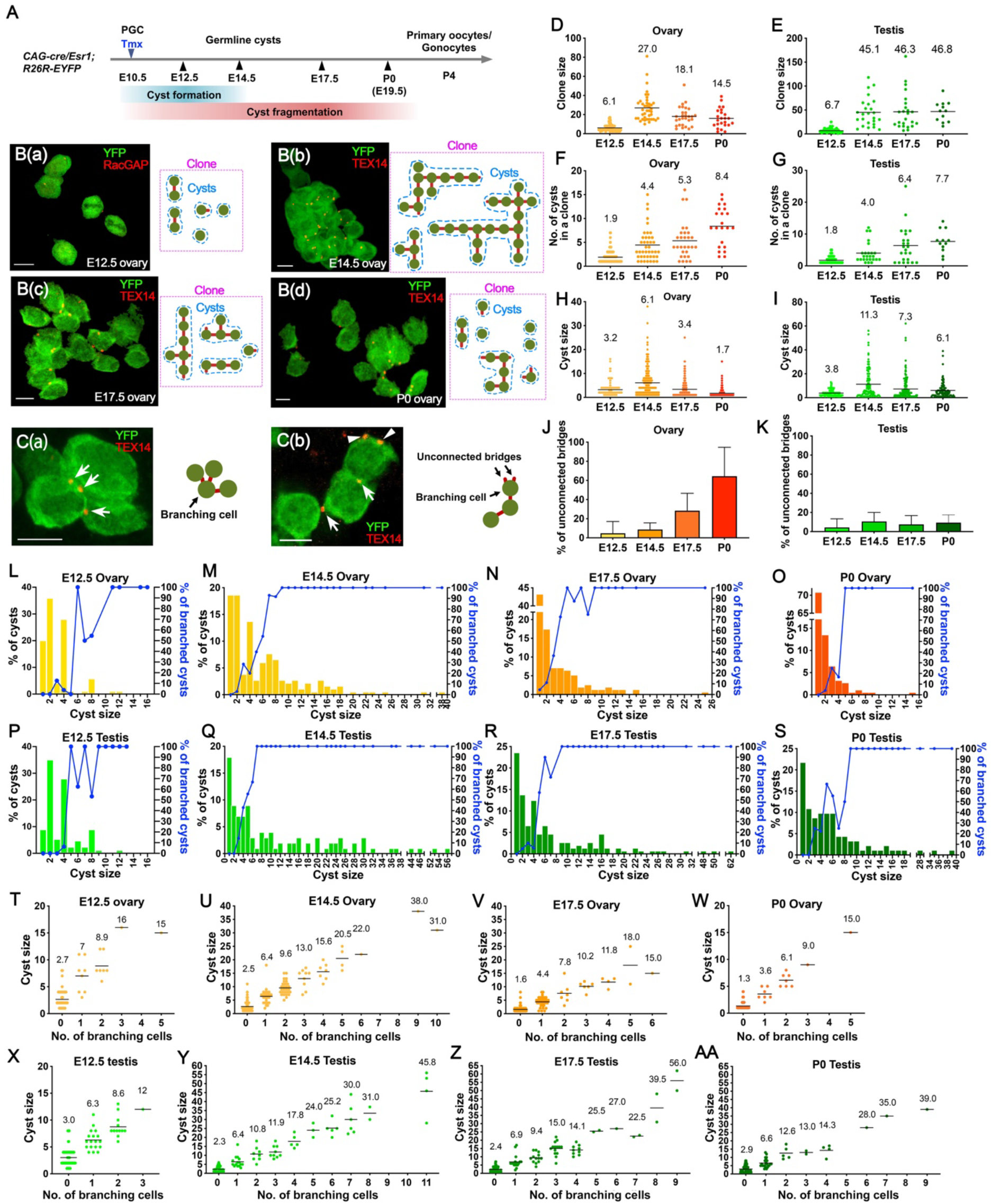
Structure of germline cysts in oocyte and gonocyte differentiation in mouse fetal gonads. (A) Time course of oocyte and gonocyte differentiation in mouse fetal gonads. A single low dose of Tamoxifen (Tmx) was injected to *CAG-cre/Esr1: R26R-EYFP* mice at E10.5 to label one primordial germ cell (PGC) per gonad on average. Fetal gonads were collected at E12.5, E14.5, E17.5 and P0 for cyst structure analysis. (B) Z-stack confocal images of lineage-labeled germ cell clones (left) and the cyst structure of the clone (right) showing the geometry of YFP-positive germ cells (green circle) and TEX14(or RacGAP)-positive bridges (red line). The cyst outlined by the blue dotted line was defined as sister germ cells that are connected by intercellular bridges. Each single germ cell was counted as a cyst. The clone outlined by the magenta dotted line was defined as sister germ cells derived from a single PGC labelled by lineage marker YFP. (C) Z-stack confocal images of lineage-labeled cysts showing an example of a branching cell (C (a)) and an example of unconnected bridges (C(b)). (D-E) Size of germ cell clones in fetal ovaries (D) and testes (E). (F-G) Number of germline cysts in a clone in fetal ovaries (F) and testes (G). (H-I) Size of germline cysts in fetal ovaries (H) and testes (I). (J-K) Percentage of unconnected bridges in fetal ovaries (J) and testes (K). (L-S) Percentage of the cysts profiled by the size of cysts (left Y axis) and percentage of the cysts that are in the branched structure (right Y axis) in fetal ovaries at E12.5 (L), E14.5 (M), E17.5 (N), P0 (O), and fetal testes at E12.5 (P), E14.5 (Q), E17.5 (R), P0 (S). (T-AA) Germline cysts profiled by cyst size and the number of branching germ cells contained in fetal ovaries at E12.5 (T), E14.5 (U), E17.5 (V), P0 (W), and fetal testes at E12.5 (X), E14.5 (Y), E17.5 (Z), P0 (AA). Scale bar=10 μm. The bar in the graph represents the average value. Data are presented as mean ± SD.

On average, each female PGC gave rise to a 27.0-cell clone at E14.5 that decreased to a 14.5-cell clone by P0 (Fig. 1D). By contrast, the clone size in fetal testes remained at ~45 cells per clone from E14.5 to P0 (Fig. 1E). This observation, at the single-germ cell-clone resolution, is consistent with previous reports of large-scale germ cell loss takes place in fetal ovaries due to oocyte differentiation but not in testes (11, 13, 31–34). From E14.5 to P0, the number of cysts per clone increased from 4.4 to 8.4 per clone in fetal ovaries, with cyst size decreasing from an average of 6.1 cells to 1.7 cells per cyst, suggesting that cyst fragmentation and cell loss take place in parallel in female cysts (Fig. 1F and H). In fetal testes, on average a E14.5 clone was comprised of 4.0 cysts, each containing 11.3 cells. By P0, cysts fragmented into 7.7 cysts, each containing 6.1 cells, suggesting that, on average, each male fetal cyst fragments once without considerable cell death (Fig 1G and I). Consequently, in P0 testes, a large number of germ cells remained connected via elongated cell protrusions with TEX14-positive bridges between the germ cells (Supplementary Fig 1). These data indicate that female and male PGCs form germline cyst from E10.5 to E14.5, despite the onset of sex differentiation around E11.5. Female cysts undergo greater fragmentation than male cysts, with germ cell loss taking place in fragmented cysts.

To detail the germline cyst structure, the geometry of the germ cells (identified by YFP) and intercellular bridges (identified by RacGAP in E12.5 goads or TEX14) in each cyst was laid out based on confocal images of individual germ cell clones (Fig. 1B). In addition to the intercellular bridge that connects two cyst germ cells, we observed unconnected bridges, which may represent the remains of the intercellular bridges following cytokinesis (Fig 1C (b))(35). From E14.5 to P0, the percentage of unconnected bridges increased from 8.7% to 61.5% of the total bridges in fetal ovaries, indicating considerable levels of cyst fragmentation occurring in female cysts (Fig 1J). By contrast, in male, the frequency of unconnected bridges remained at ~10%, consistent with lower levels of cyst fragmentation (Fig 1G, I, K).

We found that both female and male germline cysts are in branched structures, such that some germ cells are connected by three or four bridges, herein referred to as ‘branching cells’ (Fig 1C(a)). Since germline cysts form through incomplete cytokinesis during germ cell mitosis, the number of bridges reflects the rounds of division the cells have undergone. By nature, an 8-cell cyst is the stage at which branching cells first appear. However, branching cells were observed in cysts as small as 3 cells in E12.5 ovaries (Fig 1L); and in cysts as small as 5 cells in E12.5 testes (Fig 1P), reflecting cyst fragmentation during cyst formation. Further, in E14.5 ovaries, branching germ cells were observed in a few 2-cell cysts (2.94%), many 6-cell cysts (54.5%), and all cysts that are larger than 8 cells (Fig 1M). In E14.5 testes, all cysts that are larger than 6 cells were branched (Fig 1Q). Both female and male cysts retained their branched structure up until P0, with female cysts having smaller size due to extensive germ cell loss and cyst fragmentation (Fig 1N, R, O, S). We further found a linear association between the number of branching germ cells and cyst size. On average, there was one branching germ cell per 4.3 germ cells in female cysts and per 4.6 germ cells in male cysts at E14.5 when germ cell mitosis ceases, indicating that branched cyst formation may follow a similar pattern in fetal ovaries and testes. The branching germ cells persisted in both P0 fetal ovaries and testes (Fig 1T-AA and Supplementary Table 9).

### Adult male germ cells form branched cysts during spermatogenesis

During spermatogenesis in adult testes, spermatogonia stem cells (SSCs), including singly isolated cells and short cysts continually give rise to differentiating cells that form large germline cysts after a series of mitosis. Thus, we questioned whether there is difference in cyst structure between male fetal and adult cysts. We conducted single-cell lineage tracing in the adult testis to label individual SSCs using *Pax7-creER; R26R-YFP* mice (36). A single low dose of tamoxifen was injected into adult males (8-12 weeks old) and testes were collected at 1 day, 10 days and 20 days after the injection (Fig. 2A). Germ cell differentiation from SSCs is divided into three major stages based on differentiation marker expression: (1) undifferentiated spermatogonia (E-cadherin-positive/KIT-negative); (2) differentiating spermatogonia (KIT-positive); and (3) spermatocytes (synaptonemal complex protein 3, SCP3-positive). Germ cell connectivity at stage 1 and 2 involves cell membrane connections between YFP-positive sister germ cells stained with E-cadherin or KIT antibodies (Fig. 2B, D, E). TEX14-positive bridges were observed between lineage-labeled germ cells connected by E-cadherin- or KIT-positive cell membranes and between SCP3+ spermatocytes (Supplementary Fig 2A, Fig 2G).

**Figure 2.**
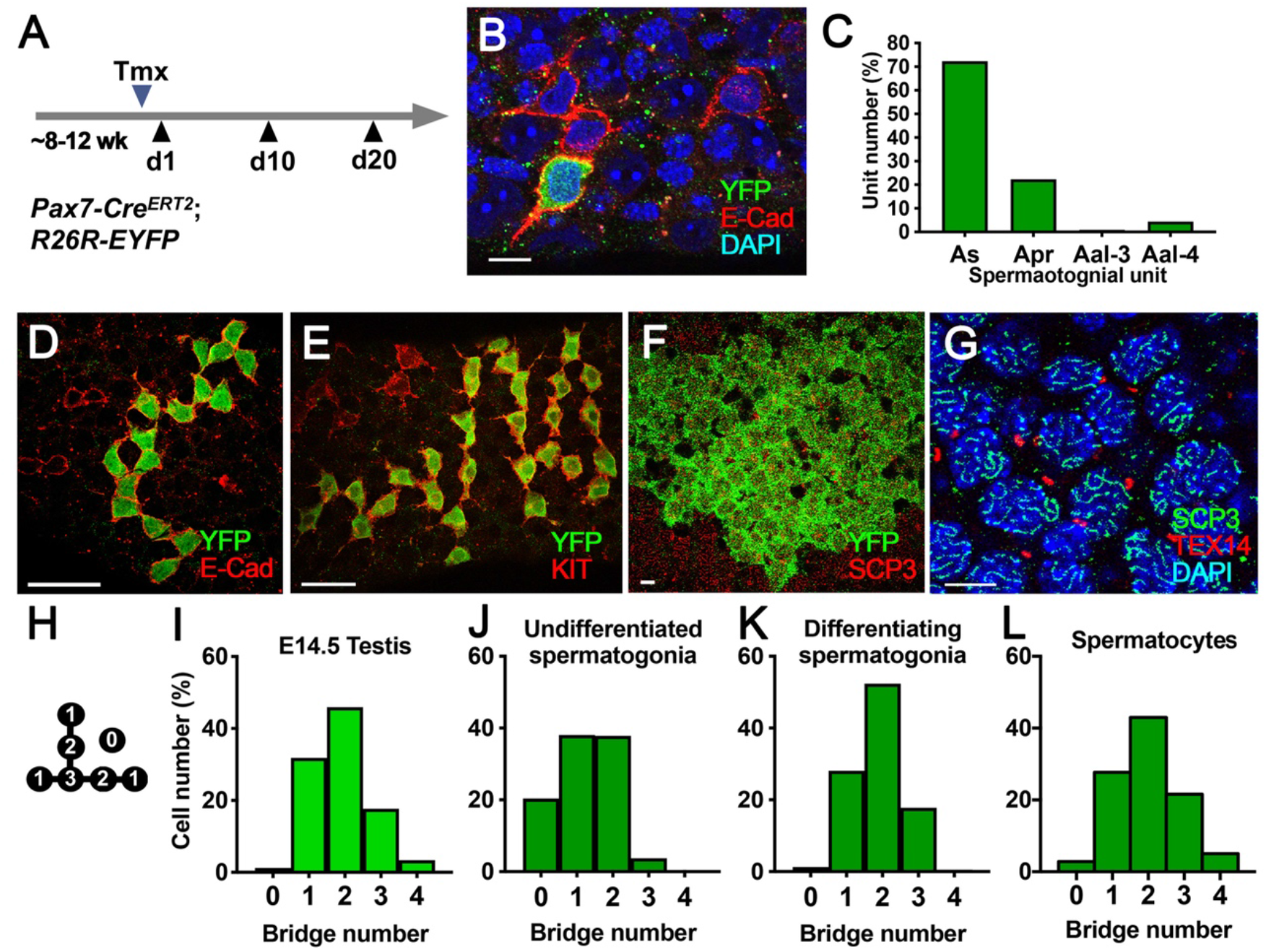
Structures of germline cysts in spermatogenesis in the adult testis. (A) Diagram showing the strategy for lineage-labeling individual Pax7-positive spermatogonia stem cells (SSCs) in the adult testis. Testes were collected on day1, 10 and 20 after Tamoxifen (Tmx) injection. (B) Confocal image showing a lineage-labelled YFP-positive undifferentiated spermatogonia expressing E-cadherin (red) one day after Tmx injection. (C) Composition of lineage-labeled spermatogonia containing different numbers of germ cells one day after Tmx injection. As: single SSC; Apr: two connected SSCs; Aal-3: three aligned SSC; Aal-4: four aligned SSCs. (D-F) Confocal images showing lineage-labeled germ cell clones that are E-cadherin positive (D), KIT positive (E) or SCP-3 positive (F) observed in the testes 10 days after Tmx injection. (G) Confocal image showing meiotic germ cells labeled by a SPC3 antibody and intercellular bridges labeled by a TEX14 antibody. (H-K) Structure of germline cysts profiled by the percentage of germ cells connected by 0, 1, 2, 3, and 4 bridges in E14.5 testes (H), E-cadherin-positive undifferentiated spermatogonia (I), KIT-positive differentiating spermatogonia (J) and spermatocytes (K). 1128 cells from 25 clones, 722 from 118 clones, 798 from 22 clones, and 640 cells were analyzed in (I), (J), (K), and (L) respectability. Scale bar=10 μm.

At 1 day after tamoxifen injection, individual linage-labeled SSCs were recognized by YFP and E-cadherin double-positive staining (Fig. 2B). Among these cells, 72.32% were single (As), 22.32% were paired (Apr; two connected SSCs), 0.89% were at Aal-3 (three aligned SSCs), and 4.46% were at Aal-4 (four aligned SSCs) (Fig. 2C). By ten days after tamoxifen injection, 92% of the labeled cysts were E-cadherin-positive/KIT-negative undifferentiated spermatogonia (Supplementary Fig 2B). To detail how cyst structure develops as spermatogenesis progresses, we quantified the number of germ cells harboring different numbers of membrane connections or TEX14-positive bridges in the clone. 0 bridge represents single germ cell. 1-bridge, 2-bridge, and 3-or 4-bridge represent the germ cell that are connected by one bridge (periphery cell), two bridges (intermediate cell), and three or four bridges (branching germ cell) respectively (Fig 2H). Given that lineage-labeled spermatocytes form several hundreds to thousands of cells, cyst structure was analyzed by counting the number of TEX14-positive bridges for each SCP-positive spermatocyte. We found that SSCs at the undifferentiated stage (E-cad-positive) were mostly connected via the linear structure, with only 3.74% of the germ cells having 3 bridges (Fig 2J). The percentage of germ cells with 3 or 4 bridges increased as spermatogenesis progressed: 18.29% in differentiating spermatogonia (KIT-positive) and 26.56% in spermatocytes (SCP3-positive) (Fig 2K). The proportion of spermatocytes with different numbers of bridges were similar with those in fetal male cysts, suggesting that male germ cells are connected in a similar configuration in both fetal and adult cysts (Fig 2L, I).

### Female and male fetal germline cysts fragment differently

The cyst number per clone and cyst size revealed that female cysts fragment to a greater extent than male cysts (Fig 1D-I). To elucidate the pattern of cyst fragmentation, we profiled the total number of bridges in each germ cell clone (Supplementary Fig 3). From E14.5 to E17.5, a female germ cell clone lost, on average, 8.9 germ cells and 9.9 connected bridges, but cyst number only increased slightly from 4.4 to 5.3, suggesting that the loss of germ cells and bridges do not cause significant cyst fragmentation during this time. From E17.5 to P0, each female germ cell clone lost, on average, 3.6 germ cells and 7.0 connected bridges, and cyst number increased from 5.3 to 8.4, suggesting that cyst fragmentation during this time may be primarily due to bridge loss (Fig 1F and Supplementary Fig 3A). On average, each male clone lost 2.2 bridges, causing limited cyst fragmentation, consistent with the presence of twice as many cysts (Fig 1G, I and Supplementary Fig 3B).

### Branching germ cells are protected in female cysts

We found that as the female cysts decreased in size from E14.5 to P0, the percentage of branched cysts and number of branching germ cells per cyst increased (Fig 3A, C). By contrast, in fetal testes, the percentage of branched cysts and number of branching germ cells per cyst remained highly similar during gonocyte differentiation from E14.5 to P0 (Fig 3B, D). These results indicate that branching germ cells are preferentially protected from cyst fragmentation and germ cell death that take place in female cysts.

**Figure 3.**
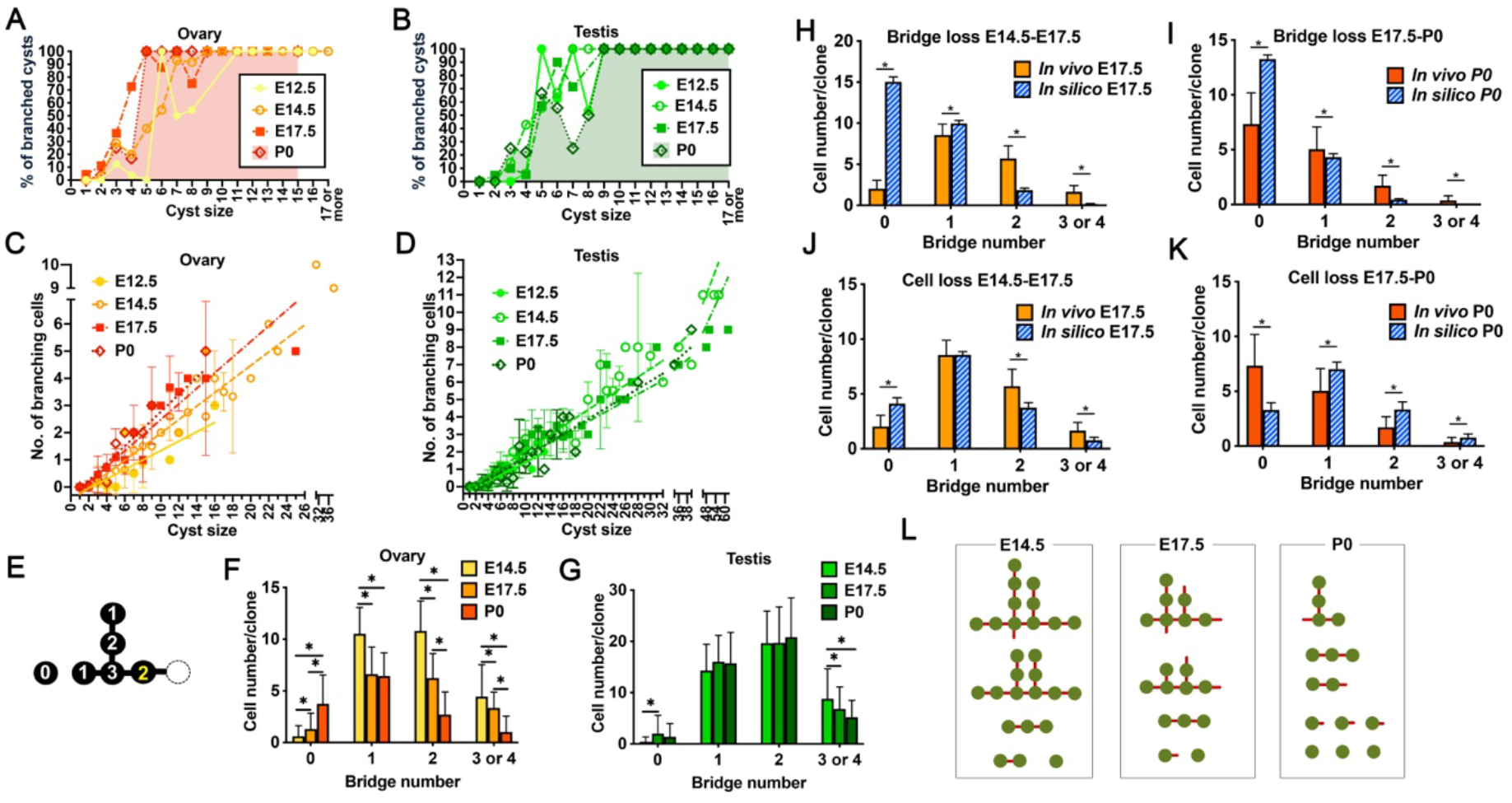
Patterns of germline cyst fragmentation due to the loss of germ cells and intercellular bridges. (A-B) Comparative profiling on the percentage of branched cysts from E12.5 to P0 in ovaries (A) and testes (B). (C-D) Comparative profiling on the number of branching cells from E12.5 to P0 in ovaries (C) and testes (D). (E-G) Changes in the number of cells that are connected with 0 bridge, 1 bridge, 2 bridges, 3 and 4 bridges in a germ cell clone in fetal ovaries (F) and testes (G) from E14.5 to P0. (H-I) Mathematic modeling of random loss of intercellular bridges in each germ cell clone in ovaries from E14.5 to E17.5 (H) and E17.5 to P0 (I). (J-K) Mathematic modeling of random loss of germ cells in each germ cell clone in ovaries from E14.5 to E17.5 (J) and E17.5 to P0 (K). (L) Proposed model of the change in cyst structure in a germ cell clone during oocyte differentiation from E14.5 to P0. The average numbers of germ cells per clone, cysts per clone, cells per cyst, bridges per clone and bridges per cyst from E14.5, E17.5 and P0 clones were used for making the model. Data in the graph are presented as mean ± SD. One way ANOVA was used for the statistical analysis between E14.5, E17.5 and P0. For modeling analyses, a two-sample t-test with 0.05-significance level was used to test the null hypothesis that the data came from independent random samples from normal distributions with equal means (*In silico*) and equal but unknown variances (*In vivo*).

To determine the location of the germ cells and bridges that are lost in cyst during oocyte differentiation, we analyzed the change in cyst structure by quantifying the number of germ cells connected by 0 bridge (single cell), 1 bridge, 2 bridges and 3 or 4 bridges (branching cell) in each germ cell clone (Fig 3E). From E14.5 to E17.5, a female clone lost, on average, 8.9 cells, which include 3.9 1-bridge cells, 4.6 2-bridge cells, and 1.1 3- or 4-bridge cells. These cells are 37% of the total 1-bridge cells, 42% of the 2-bridge cells, and 25% of the 3- or 4-bridge cells of a clone, respectively (Fig 3F). From E17.5 to P0, a female clone lost on average, 3.6 cells, which include 0.2 1-bridge cell, 3.5 2-bridge cells, and 2.3 3- or 4-bridge cells, i.e. 3% of the total 1-bridge cells, 56% of the 2-bridge cells and 70% of the 3 or 4-bridge cells of a clone, respectively (Fig 3G). As cyst fragmentation progresses, the number of bridges connected to the germ cell decreases, i.e. 3- or 4-bridge germ cells become 0-, 1-, or 2-bridge germ cells. Our results show that although 1- or 2-bridge cells occur frequently during cyst fragmentation, the majority of cell loss involved 1- and 2-bridge cells during E14.5 to E17.5. Our results suggest that germ cell loss in cysts from E14.5 to E17.5 takes place in a selective manner with 1-and 2-bridge cells being lost and branching germ cells being protected preferentially.

In male germ cell clones, the germ cells connected by 1-or 2-bridges remained approximately the same; however, on average, the clone lost 3.6 branching germ cells from E14.5 to P0, indicating that the limited cyst fragmentation in male cysts may preferentially take place at branching germ cells (Fig 3G).

To test our hypothesis that cyst fragmentation and germ cell loss in female cysts occur in a nonrandom manner with branching cells being protected, we mathematically modelled random bridge loss and germ cell loss in each female germ cell clone and compared the resultant cyst structure from *in silico* modeling with the observed *in vivo* cyst structure (Fig 3H-K). With mathematic modeling considering connected intercellular bridges only, unconnected bridges were excluded when counting the number of bridges connected to each *in vivo* germ cell in order to make a comparison between *in silico* and *in vivo* cyst structure. Based on the change in average bridge number per clone from E14.5 to E17.5, 44% of the bridges were randomly removed from each E14.5 female clone (*in vivo* E14.5, n=41) and the cyst structures generated by modeling (*in silico* E17.5) were compared with the actual cyst structure found *in vivo* in E17.5 clones. Similarly, 55% of the bridges were randomly removed from each E17.5 female clone (n=29), and the cyst structures generated by modeling (*in silico* P0) were compared with actual cysts found in the *in vivo* P0 clones. This modeling, in which the total cell numbers do not change, demonstrated that random removal of the bridges would generate significantly more 0-bridge single cells than observed *in vivo*. Moreover, fewer 2-bridge and 3- or 4-bridge cells appeared in modeled clones compared with those *in vivo*, suggesting that cyst fragmentation does not take place in a random manner, instead it occurs with bridges connected to branching germ cells being protected (Fig 3H, I).

We further modeled random germ cell loss (Fig 3J and K). On average, 33% of the germ cells were lost in a clone from E14.5 to E17.5 and 20% of the germ cells were lost in a clone from E17.5 to P0 *in vivo* (Fig 1D). When germ cells (and the associated bridges) were removed randomly at such rates in each E14.5 germ cell clone (n=41), there were significantly fewer 2-bridge and 3- or 4-bridge germ cells in *in silico* E17.5 clones than *in vivo*, suggesting that branching germ cells are preferentially protected *in vivo* during E14.5 to E17.5. In contrast, when 20% of the germ cells were randomly removed from E17.5 clones (n=29), significantly more germ cells remained connected in *in silico* P0 clone, suggesting extensive cyst fragmentation occurs from E17.5 to P0 *in vivo*.

Based on the averaged cell number per clone and per cyst, as well as germ cells connected with 1, 2, and 3 or 4 bridges per clone and per cyst, the averaged change in cyst structure was modeled in Fig 3L. Together with mathematic modeling analyses, our results suggest that from E14.5 to E17.5, germ cells are lost in a selective manner, in which branching germ cells and the associated bridges are preferentially protected and periphery germ cells are lost; and from E17.5 to P0, cysts undergo extensive fragmentation, primarily due to bridge loss, breaking down into small cysts and individual germ cells (Fig 3L).

### Female branching germ cells contain enriched cytoplasm and organelles

Our previous and present studies demonstrated that organelle and cytoplasmic transport between cyst germ cells results in two germ cell populations by P0: (1) germ cells with enriched cytoplasm and organelles, thus containing B-bodies, are preferentially protected from cell death and become primary oocytes; and (2) germ cells that are smaller in volume, without the B-body, undergo apoptosis preferentially (18). To determine whether the branching germ cells are enriched with cytoplasm and organelles in the cyst, we measured the volume of each germ cell in the clone and plotted this against the number of bridges connected to them. We separated the germ cells into two categories: (1) single cells that have separated with the cyst and thus presumably have completed organelle and cytoplasm transport (among these cells, some still have unconnected TEX14-positive bridges, which partially reflects the history of germ cell connection); and (2) germ cells in cysts that remain connected and are in the process of organelle and cytoplasm transport.

Consistent with our previous study, the volume of cyst germ cells increased as oocyte differentiation progressed from E14.5 to P0 (Supplementary Fig 4A)(18). In E14.5 germ cell clones (6.1 cells per cyst), single cells comprised 2.1% (12/564) of the total germ cells and single germ cells without bridge had the smallest volume (514 μm^3^). Branching germ cells, comprising 16.1% (91/564) of the total germ cells, had the largest volume (713 μm^3^) (Fig 4A). By E17.5 (3.4 cells per cyst) 13% (38/292) of the germ cells were single cells, of which a small fraction (13/292) had unconnected bridge remnants. The single germ cells with 2 or more unconnected bridges had a larger volume compared to single germ cells without bridges on average. Among germ cells in cysts, bridge number and cell volume showed a positive correlation, with branching germ cells having the largest volume (817 μm^3^) (Fig 4B). By P0 (1.7 cells per cyst), 41.7% (128/307) of the total germ cells were single cells, and germ cells without bridges or with single unconnected bridges had a wide range of cell volumes, indicating that by the time of cyst fragmentation is complete, germ cells have a divergent range of cytoplasmic content, due to cytoplasmic transport in germline cysts. On average, single germ cells were larger in volume compared to germ cells in cysts in P0, suggesting that germ cells in cysts have not completed cytoplasmic transport at this time (Fig 4C, Supplementary Fig 4B). Among germ cells in cysts, germ cells with 1, 2 or multiple bridges were comparable in volume, smaller than single germ cells, and of a similar cell volume to the branching germ cells in E17.5 cysts, indicating that germ cells in P0 cysts are in the later stages of cytoplasmic transport (Fig 4C).

**Figure 4.**
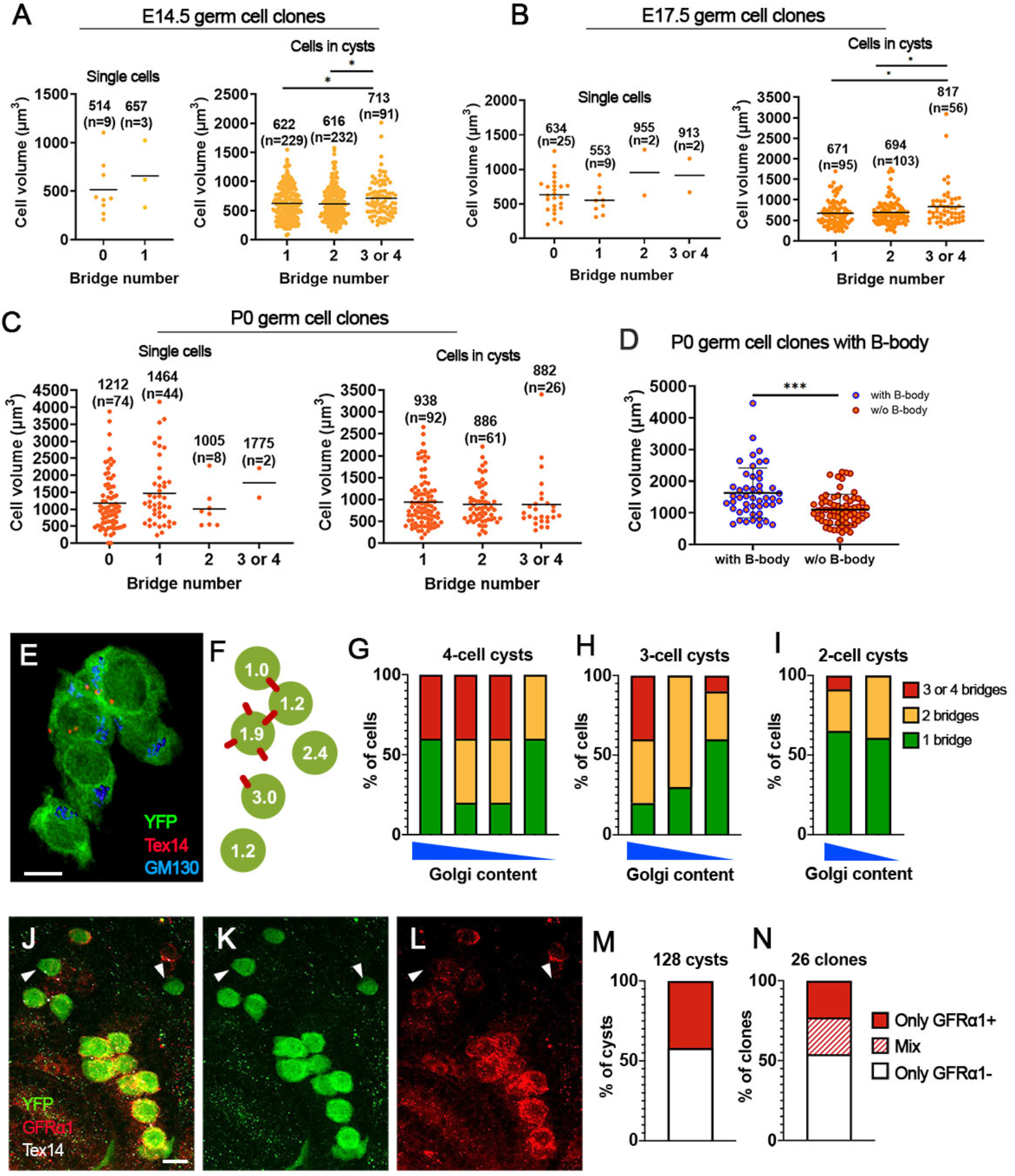
Cytoplasm and organelle content in cyst germ cells profiled based on bridge number during oocyte differentiation. (A-C) The volume of single germ cells and germ cells in cysts in E14.5, E17.5, and P0 germ cell clones. The bar in the graph represents the average value; the number above the bar represents the average volume of the germ cells; the number in the parentheses represents the number of measured germ cells. (D) Comparison of the germ cell volume between the germ cells with a B-body and without a B-body in P0 clones. Data in the graph are presented as mean ± SD and a t-test was used for statistical analysis. (E) Z-stack confocal image showing a lineage-labeled germ cell clone. Germ cells were revealed by YFP antibody staining, bridges were stained with a TEX14 antibody, and Golgi complexes were stained with a GM130 antibody. (F) Diagram of the germ cell clone in (E) showing the geometry of the germ cells (green circles) and the bridges (red lines) connected with them. The numbers in the germ cells represent the normalized value of the content of Golgi complexes in each germ cell. (G-I) Cyst germ cells were profiled based on the number of bridges connected with them and the amount of Golgi complexes each germ cell contained in 4-cell cysts (G), 3-cell cysts (H) and 2-cell cysts (I). (J-L) Confocal images showing a lineage-labeled germ cell clone in the P0 testis. The testis was stained with a GFP antibody for cyst germ cells, a TEX14 antibody for intercellular bridges and an antibody for spermatogonia stem cell marker GFRα1. Arrowheads indicate lineage-labeled germ cells with negative GFRα1 staining. (M-N) Percentage of cysts (M) and clones (N) containing the germ cells with only GFRα1 positive germ cells (red box), only GFRα1 negative germ cells (white box) and both GFRα1 positive and negative germ cells (stripped box). Scale bar=10 μm.

To determine whether branching germ cells are also enriched with organelles, we quantified the Golgi complex volume in each germ cell of individual cysts using confocal imaged germline cysts and Imaris Cell Imaging software (Fig 4E, F). We focused on 4-cell, 3-cell, and 2-cell cysts from E18.5 and P0 mouse ovaries, which are in the later stages of organelle transport in cysts. We found that the number of bridges corresponded positively with the volume of Golgi complexes in the cell (Fig 4G-I). This suggests that branching cells preferentially accumulate organelles within germline cysts. Our previous study demonstrated that in the germ cells with enriched organelles, Golgi complexes organize into a B-body and the germ cells with a B-body are preferentially protected from apoptosis in postnatal ovaries to become primary oocytes (18). To directly correlate germ cell volume, organelle enrichment, and oocyte fate among the germ cells derived from the same PGC, we analyzed germ cell volume in P0 germ cell clones containing germ cells that have completed B-body formation. In each germ cell clone, germ cells with a B-body had a larger cell volume compared to germ cells without a B-body, reinforcing that germ cells with enriched cytoplasm preferentially differentiate into primary oocytes (Fig 4D). Together, these findings suggest that branching germ cells accumulate organelles and cytoplasm from other cyst germ cells and differentiate into primary oocytes.

### Uniform SSC marker expression in male fetal germline cysts

To analyze whether fetal germ cells in branched male cysts show heterogeneity in cell fate, we examined the expression of spermatogonia stem cell (SSC) markers GFRα1 (GDNF family receptor alpha-1)(37, 38), PLZF (promyelocytic leukemia zinc finger) (39, 40) and UTF1 (undifferentiated embryonic cell transcription factor 1)(41, 42) in lineage-labeled germ cell clones in P0 testes. For GFRα1, either all cells in the cyst were positive (42% of the cysts) or all cells in the cyst were negative (58% of the cysts) regardless of cyst structure (Fig 4J-M), suggesting that cyst structure does not influence SSC differentiation in male cysts. The germ cells in a clone did not express GFRa1 in a synchronized manner, despite being derived from a single PGC. Approximately 23% of clones contained only GFRa1-positive cysts, 23% of clones contained both GFRa1-positive cysts and negative cysts, and 54% of clones contained only GFRa1-negative cysts in P0 testes (Fig 4N). PLZF showed a similar consistent expression in the germ cell in the cyst but not in the clone (Supplementary Fig 5). These results indicate that GFRa1 and PLZF expression may be regulated by signaling molecules that are shared among germ cells within a male cyst rather than intrinsically. UTF1 was expressed in all P0 germ cells, without detectable difference (Supplementary Fig 5).

### Female germ cell clusters identified by single-cell RNA-seq

To reveal whether female germ cells are different in RNA composition, we conducted single cell RNA sequencing on E14.5 and P0 ovarian cells. 18 clusters of cells were identified based on their molecular profiles, and among those, clusters 4, 5, and 13 were identified as germ cells, based on the expression of the germ cell-specific gene *Ddx4* (Fig 5A and B), comprising 53.8±0.47%, 38.7±0.27%, and 7.5±0.74% of the total E14.5 germ cells, and 46.4±0.96%, 43.0±0.48%, and 10.6±1.4% of the total P0 germ cells, respectively. We profiled total transcripts and the number of detected genes as indicators for RNA content. Germ cells in clusters 4 and 5 at E14.5 had a wide range of RNA content; on average, cluster 13 contained the highest number of transcripts and detected genes with less variation. Among P0 germ cells, cells in clusters 4 and 5, and particularly cluster 5, had diverged into two sub-groups, with the majority of the germ cells being low in numbers of transcripts and detected genes, while germ cells in cluster 13 remained the highest in RNA content on average (Fig. 5C and D).

**Figure 5.**
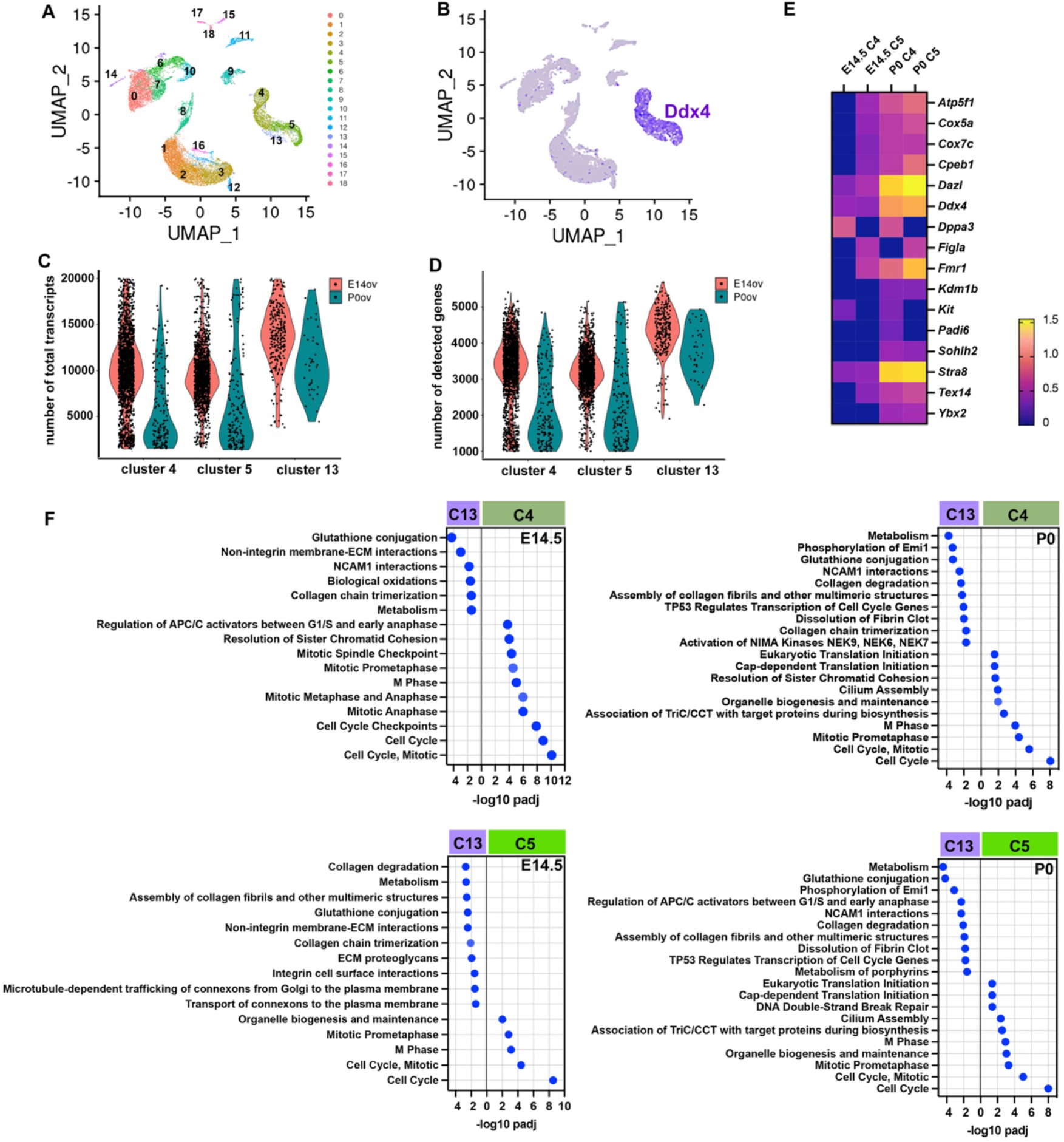
Germ cell clusters in fetal ovaries identified by single cell RNA-seq. (A) Thirteen clusters of cells identified by single cell RNA-seq in E14.5 and P0 mouse fetal ovaries. (B) Clusters 4, 5 and 13 are identified as germ cells based on their expression of germ cell-specific gene *Ddx4*. (C) Violin plot showing the distribution of the total transcripts (mRNA) in each cell in the clusters 4, 5, and 13. Each dot represents a cell. (D) Violin plot showing the detected genes in each cell in the clusters 4, 5, and 13. Each dot represents a cell. (E) Differential expression of the genes involved in oocyte differentiation in clusters 4 and 5 shown in log2 fold change compared to cluster 13. (F) REACTOM analysis on the differentially expressed genes between germ cells in clusters 4 or 5 and cluster 13 at E14.5 and P0.

Compared with clusters 4 and 5, cluster 13 germ cells had lower mRNA expression of genes that are involved in oocyte differentiation and development (*Cpeb1, Dazl, Ddx4, Dppa3, Figα, Fmr1, Kdm1b, Kit, Padi6, Sohlh2, Stra8, Ybx2*,)(43–54), mitochondrial function (*Atp5f1, Cox5a, Cox7c*)(55)(UniProt) and of the intercellular bridge protein *Tex14* (Fig 5E)(14), suggesting that cluster 13 may not represent the germ cells differentiating into primary oocytes.

To better understand the molecular features of these three clusters of germ cells, we compared the biochemical pathways of these cells at E14.5 and P0 using GO term REACTOM analysis (Supplementary Table 10-15). The differentially expressed genes between clusters 4 and 5 are cell cycle-related at both E14.5 and P0, suggesting that clusters 4 and 5 may represent germ cells at different meiotic stages (Supplementary Fig 6). At E14.5, comparison between clusters 4 and 13 revealed that cluster 4 cells are enriched in cell cycle pathways, and cluster 13 cells are enriched in genes related to metabolism, collagen chain trimerization, and extracellular matrix (ECM) organization. Comparison between clusters 5 and 13 showed that cluster 5 cells are enriched for organelle biogenesis and maintenance, and four cell cycle pathways. Cluster 13 cells are enriched for 12 pathways, which include function and organization of collagen and ECM, as well as intracellular trafficking (transport of connexons to the plasma membrane and microtubule-dependent trafficking of connexons from Golgi to the plasma membrane) (Fig 5F).

At P0, comparisons between clusters 4 and 13 revealed that cluster 4 cells are enriched for seven cell cycle-related pathways and one pathway related to protein metabolism (association of TriC/CCT with target proteins during biosynthesis), two pathways related to organelle biogenesis (organelle biogenesis and maintenance, cilium assembly), and two pathways related to translational regulation (Cap-dependent translation initiation, eukaryotic translation initiation); cluster 13 cells are enriched for 11 pathways, including cell cycle, metabolism, and ECM organization. In addition, the difference in REACTOM pathways between clusters 13 and cluster 5 is very similar to that between clusters 13 and cluster 4 (Fig 5F).

Gene ontology biological pathway analysis revealed that germ cells in cluster 13 are enriched with genes involved in the regulation of cell death and apoptotic processes when compared with cluster 5 at E14.5 and P0. Taken together, these results suggest that cluster 13 germ cells may represent intermediate germ cells during cytoplasmic transport, with a relatively higher level of ECM and cytoskeleton organization that is required for receiving and exporting cytoplasmic content, and these cells further undergo cell death after donating their cytoplasmic content. We attempted to locate cluster 13 germ cells in the germline cyst by antibody staining of the proteins that have higher expression from single cell-RNA seq results, including *Espn, Stmn1, Cald1, Ezr, S100A10. Gstm1, Gstm2, Tsc22d1, Tmsb4x* and *Tmp1*. However, we could not distinguish certain germ cell populations in the cyst (Supplementary Table 16).

### E-cadherin facilitates the formation of branched cysts

The germline cysts arising from a single PGC have a linear structure at the 4-cell stage after two rounds of synchronized mitotic divisions (Fig 6A). When the interconnected four germ cells divide synchronously into an 8-cell cyst, the following three types of cysts can be produced: (1) a linear cyst; (2) a branched cyst with one branching germ cell (1-branching cell cyst); and (3) a branched cyst with two branching germ cells (2-branching cell cyst). If branched cysts are produced randomly due to mitotic spindle orientation, the expected cyst percentage is 25%, 50%, and 25% for linear, 1-branching cell cysts, and 2-branching cell cysts respectively (Fig 6B, Supplementary Fig 7). However, when we analyzed 8-cell cysts collected from E12.5 mouse ovaries, we found that the cysts were 48% linear, 19% 1-branching cell cysts, and 33% 2-branching cell cysts, suggesting that branched cyst formation in mouse fetal ovaries is not due to random spindle orientation during germ cell mitosis.

**Figure 6.**
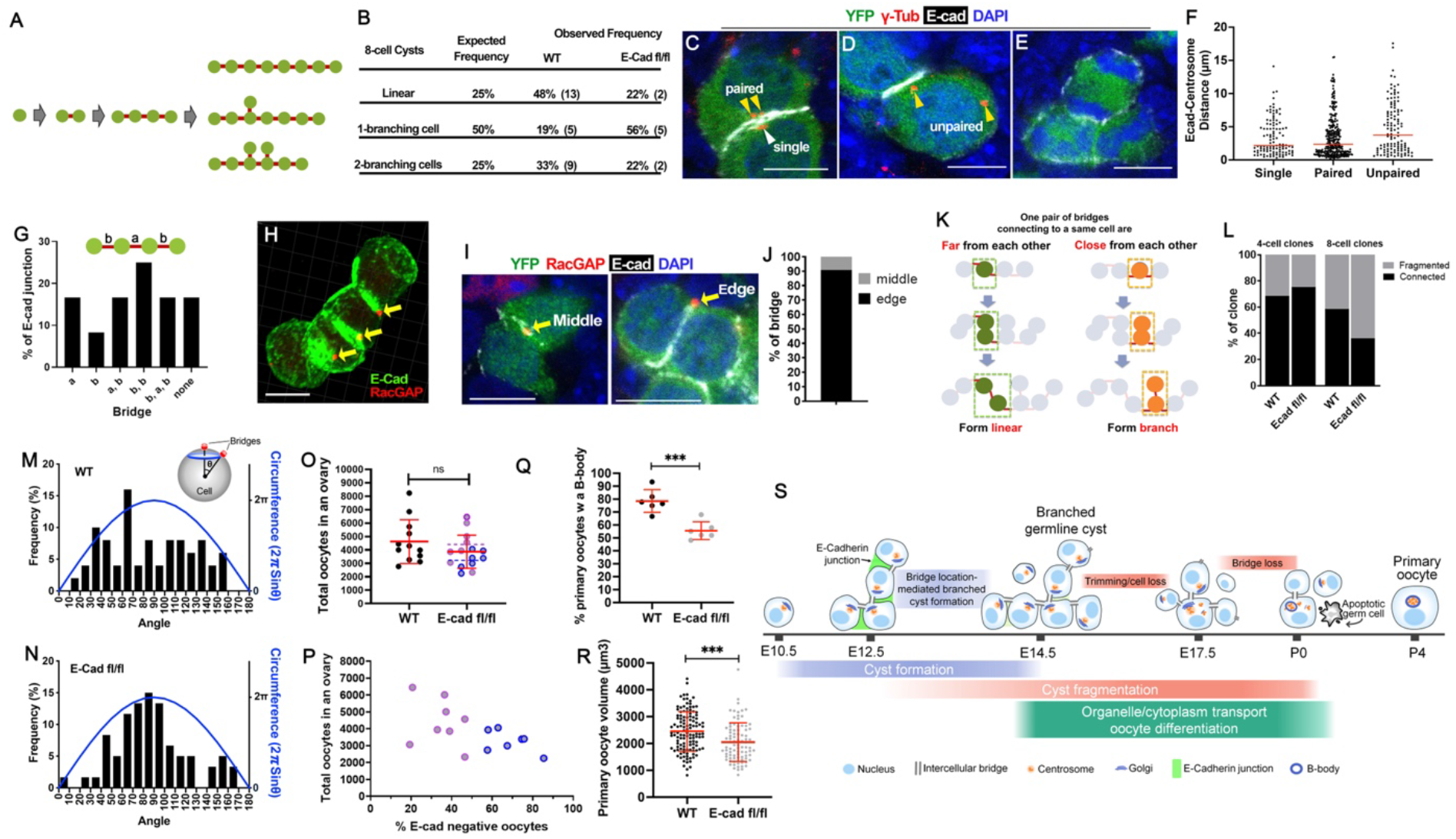
E-cadherin junctions are involved in the formation of branched cysts by positioning intercellular bridges. (A) Diagram demonstrating three types of 8-cell cysts (linear, 1-branching cell and 2-branching cell) that form through incomplete cytokinesis after synchronized mitotic division of a 4-cell cyst. (B) Percentage of three types of 8-cell cysts expected from random spindle orientation and observed in wildtype (WT) and E-cadherin knockout (E-cad fl/fl) germ cell clones. (C-D) Confocal images showing centrosome locations (labeled by γ-tub antibody staining) and the E-cadherin junctions in non-dividing cyst germ cells. Yellow arrowheads in C indicate two paired centrosomes. White arrowhead in C indicates a single centrosome. Yellow arrowheads in D indicate two unpaired centrosomes. (E) Confocal image showing that in a cyst with mitotic germ cells, E-cadherin was found along the cell membrane without a junction between germ cells. (F) The distance between E-cadherin junction and centrosomes in 4-cell germline cysts. (G) The distribution of E-cadherin junctions in 4-cell cysts. a, b and c represent the bridges between two germ cells. (H) 3-D reconstructed confocal image showing the location of E-cadherin junctions and intercellular bridges stained by a RacGAP antibody. (I) Confocal images showing an intercellular bridge found in the middle and at the edge of the E-cadherin junction. (J) Percentage of the intercellular bridges located in the middle or at the edge of the E-cadherin junction in 4-cell cysts. (K) Diagram showing that the location of intercellular bridges influences the structure of the branched cyst during cyst formation. (L) The percentage of fragmented and connected cysts in the wildtype and E-cadherin knockout 4-cell and 8-cell cysts. (M, N) Distribution of the angles measured between two adjacent intercellular bridges in the wildtype (M) and E-cadherin knockout (N) 4-cell cysts. (O, P) Number of total oocytes in the P4 wildtype and E-cadherin knockout ovaries. Graph in (P) showing a negative association between the percentage of E-cadherin-negative oocytes and the number of oocytes in the ovary. Magenta-lined dots: ovaries with less than 50% E-cadherin-negative oocytes; Blue-lined dots: ovaries with more than 50% E-cadherin-negative oocytes. (Q) Percentage of the primary oocytes containing a B-body in the P4 wildtype and E-cadherin knockout ovaries. (R) Volume of primary oocytes in the P4 wildtype and E-cadherin knockout ovaries. (S) A diagram demonstrating the timeline of cyst formation, fragmentation and cytoplasm and organelle transport-mediated oocyte differentiation in germline cysts. Scale bar=10 μm. Data in the graph are presented as mean ± SD. The bar in the graph represents the average value. T-tests were used for the statistical analysis between the two groups.

In many cell types, including germline stem cells in *Drosophila* testes, cadherin junctions formed between the cells orient mitotic spindles during division (56, 57). We examined the distribution of centrosomes and E-cadherin in 4-cell cysts. E-cadherin junctions were found in 4-cell cysts between two adjacent cyst germ cells, without a preference in forming between certain two germ cells (Fig 6C, G). Germ cell centrosomes tended to localize near the E-cadherin junctions in interphase germ cells (Fig 6C, D). Centrosome location may not be associated with the cell cycle given that the distance between the centrosome and E-cadherin junctions varied from less than 1 μm to over 10 μm for single centrosomes, duplicated centrosomes (paired), and two separated centrosomes (unpaired) (Fig 6C-F). Despite the association between centrosomes and E-cadherin junctions, the junctions disappeared when germ cells enter the M-phase, suggesting that E-cadherin junctions may not be involved in spindle orientation during cyst formation (Fig 6E, Supplementary Movie 1).

We found that intercellular bridges in cysts were located at the edge of E-cadherin junctions at a high frequency (Fig 6H, I). In 4-cell cysts, 90.7% of the bridges were found at the edge of the junction (Fig 6J). The location of the bridges influences the location of future daughter cells and branched cyst structure as demonstrated in Fig 6K and Supplementary Fig 7. To elucidate whether E-cadherin junctions are involved in positioning bridges between germ cells and in branched cyst formation, we knocked out E-cadherin in lineage-labeled individual PGCs. We compared cyst formation and structure between mutant and wildtype cysts using *CAG-creER+/-; R26R-YFP/YFP; E-cadherin-flox/flox* mice, where the injection of a low dose of tamoxifen induces cre recombinase activity that drives the expression of YFP protein and depletion of E-cadherin simultaneously in the single PGCs. E-cadherin knockout led to increased cyst fragmentations at the 8-cell cyst stage (Fig 6L), indicating that E-cadherin junctions are involved in holding cyst germ cells together, in combination with the intercellular bridges during cyst formation. We measured the angles generated from the center of the cell to each of the two bridges that it has in 4-cell cysts and found that they were distributed over a wide range of values in 4-cell wild-type cysts. By contrast, angles measured in the E-cadherin knockout 4-cell cysts had a distribution with the majority being ~90 degrees, showing a close coincidence with the pattern of random bridge distribution on the cell surface based on calculated circumferences (Fig 6M, N). The change in bridge location at the 4-cell stage led to an altered percentage in 8-cell cysts of three types of structures. The percentages of linear, 1-branching cell cysts, and 2-branching cell cysts were 22%, 56% and 22%, respectively, which is very similar to the expected ratio when 8-cell cysts form randomly (Fig 6B, Supplementary Fig 7).

To investigate the effect of altered cyst structure and cyst fragmentation on oocyte differentiation, we examined oocyte differentiation in *Figα-cre+/-; E-cadherin-flox/flox* mice where E-cadherin knockout occurs specifically in germ cells, starting at around E12.5 (58). We found that the efficiency of E-cadherin knockout in germ cells (i.e. percentage of E-cadherin negative oocytes) varied in the mutant ovaries and no significant difference was observed in average number of oocytes per ovary between wildtype and mutant ovaries at P4 (Fig 6O). However, there was a strong negative association between the number of oocytes in the ovary and the percentage of E-cadherin negative oocytes, suggesting that the loss of E-cadherin causes reduced oocyte numbers (Fig 6P). Primary oocytes with E-cadherin knockout were smaller in cell volume and showed a reduction in the percentage of primary oocytes containing a B-body, indicating that altered branched cyst formation and increased cyst fragmentation in E-cadherin mutant germ cells lead to defects in organelle and cytoplasmic enrichment during oocyte differentiation (Fig 6Q, R).

## Discussion

### Branched cyst structure maximizes intercellular communication

During gametogenesis in germline cysts, intercellular bridges allow the exchange of signaling molecules and small organelles between cyst germ cells. Germline cysts in many animals are in branched structures, including *Drosophila* ovaries and testes, *Xenopus* ovaries, and rat adult testes (19) (59, 60). The pattern of physical connections between germ cells likely influences intercellular communications. Here, we have found that germline cysts in mouse fetal ovaries and testes are in branched cyst structures, with an average of 16.8% female germ cells and 20.4% male germ cells being connected by three or four intercellular bridges by the time cyst formation ceases at E14.5. Similarly, during spermatogenesis in adult testes, cysts become more branched as spermatogenesis progresses such that 26.6% of the germ cells are branching cells when they enter meiosis.

The branched cyst structure may provide an advantage in facilitating germ cell communication, compared with linear cysts, given some germ cells are connected with more than two bridges. To analyze the difference in the efficiency of germ cell communication between different cyst structures, we compared the efficiency of information sharing in three types of 8-cell cysts using mathematic modeling, which measured the time needed to spread the information from a single cell until it reaches a steady-state distribution among the cells in the entire 8-cell cyst (Supplementary file 1). The distribution time in the linear cyst, one-branching cell cyst, and two-branching cell cyst were 40 sec, 30 sec and 22 sec, respectively, thus branched cyst structures facilitate more efficient intercellular communications, compared with linear cysts (Supplementary Movie 2-4). This result may explain why male germ cells form branched cyst structures without significant fragmentation to ensure efficient exchange of signaling molecules needed for gonocyte differentiation and spermatogenesis.

### Selective loss of germ cells and intercellular bridges facilitate oocyte determination

In mouse germline cysts, cyst fragmentation due to bridge loss and germ cell loss takes place in female cysts largely during oocyte differentiation. Our present study revealed that female cysts fragment differently at two stages of oocyte differentiation in fetal ovaries. From E14.5 to E17.5, 9 germ cells were lost from 4.4 cysts per E14.5 clone and resulted in 5.3 cysts per E17.5 clone, indicating that germ cell loss results in decreased cyst size without causing significant cyst fragmentation during this time; and that the cell loss is primarily due to the loss of cells connected by 1 or 2 bridges within the cyst (Fig. 3F). E17.5 clones contained very few single cells, this indicates that either single cells are eliminated immediately after being fragmented from E14.5 cysts; or eliminated after cyst germ cells form syncytia due to fragmented cell membrane observed in wildtype and *Tex14* mutant fetal ovaries (16, 18). The selective germ cell loss within the cyst from E14.5 to E17.5 may help facilitate the directional communication among cyst germ cells, which is needed for organelle and cytoplasmic enrichment as cysts undergo rapid fragmentation from E17.5 to P0, caused primarily by bridge loss. It is worth noting that, on average, 5.3 germline cysts are found in E17.5 ovaries and 6 primary oocytes are found in P4 ovaries (Fig 1F) (11). Whether each E17.5 cyst has a well-established intercellular polarity that facilitates directional organelle and cytoplasmic transport into specific germ cells from E17.5 to P0 will be investigated in our future studies. Mechanisms for selective bridge closure/loss between E17.5 and P0, in particular, whether the cellular machinery of cytokinesis is found at certain bridges, should be addressed after acquiring a better understanding of bridge composition and closure.

The number of bridges on the germ cell reflects the rounds of mitosis the cell has experienced. The branching germ cells connected with three or four bridges are likely the initial founder cells of the cyst, since, on average, each PGC undergoes five rounds of mitotic division to give rise to a 27-cell clone. We found that from E14.5 to E17.5, branching germ cells are preferentially protected from cell death and cyst fragmentation, increasing their chances of enriching their cytoplasm and becoming oocytes. How branching germ cells in mouse germline cysts are prioritized for organelle accumulation remains unclear. Our analyses on the change in cyst structure suggest that cytoplasmic transport in germline cysts is conducted in a directional manner. From E14.5 to E17.5, the preferential loss of 1-bridge and 2-bridge germ cells indicates that cellular content is first transported locally between peripheral germ cells and further enriched in branching germ cells. The distribution of proteins that are involved in cellular polarity and cytoskeleton organization in cyst germ cells will be investigated in our future study. We found that germ cells had a comparable cell volume in the cysts at the late stage of transport in P0 ovaries, in contrast to the different amounts of Golgi content in the cyst germ cells at this stage (Fig 4C and G-I). This result suggests that the Golgi complex may be transported before the rest of the cytoplasmic content; and as the organizing center of microtubules, Golgi complexes and the associated centrosomes may be involved in mediating the direction of organelle and cytoplasm transport between cyst germ cells.

### E-cadherin junctions facilitate branched cyst formation

We observed E-cadherin junctions between cyst germ cells during cyst formation from E10.5 to E14.5, and the junctions diminished after E14.5, suggesting that they may be involved in germ cell mitotic division (Supplementary Fig 8). The E-cadherin junctions may be essential for holding cyst germ cells together, which are otherwise connected via intercellular bridges. E-cadherin junctions thus may protect cysts from fragmentation caused by somatic cell invasion during cyst formation, an idea confirmed by our observation that E-cadherin-depleted germ cell clones had a higher rate of cyst fragmentation (Fig 6L).

We found that E-cadherin junctions play a role in branched cyst formation through positioning the intercellular bridge between two sister germ cells (Fig 6K). E-cadherin is involved in spindle orientation in *Drosophila* germline stem cells, as well as sensory organ precursor cells and epithelial cells in *Drosophila* and mammals (22, 61–67). Here, we found that, although germ cell centrosomes showed a strong association with the E-cadherin junctions during interphase, the junctions became dispersed during mitotic division, indicating that E-cadherin junctions in the germline cyst are not involved in orienting mitotic spindles during germ cell division. It is intriguing that when E-cadherin was depleted in individual 4-cell cysts, both the angles between two adjacent bridges and the percentages of linear and branched 8-cell cysts resembled the pattens caused by random positioning of intercellular bridges by E-cadherin junctions (Fig 6B, N). This result indicates that branched cyst formation in wildtype ovaries is not caused by randomly positioned intercellular bridges. The dynamics of E-cadherin junction formation and maintenance between cyst germ cells may provide insight into how bridge location and resultant branched cysts formation are regulated.

We observed that in mice with germ cell-specific depletion of E-cadherin, mutant primary oocytes had a decreased cell volume and the percentage of primary oocytes containing a B-body, suggesting that cytoplasmic transport in these germ cells is compromised. However, given the role of E-cadherin in PGC survival and cell polarity, how altered cyst structure changes oocyte determination in germline cysts needs to be addressed in future studies with the appropriate genetic mutant mouse models.

### Oocyte is primarily determined by cytoplasmic transport

Our single cell RNA-seq data identified three germ cell clusters, with cluster 13 germ cells being the least abundant at both E14.5 (beginning of the organelle transport) and P0 (end of the organelle transport), but having, on average, the highest level of RNA content, as indicated by the number of transcripts and detected genes. Using single-PGC lineage tracing and total germ cell counts, our previous study found that ~20% of the E14.5 germ cells and ~40% of the P0 germ cells differentiate into primary oocytes by P4 (11). In the present study, we found that 7.5% of germ cells fell into cluster 13 at E14.5 and 10.6% fell into cluster 13 at P0. Cluster 13 germ cells also had lower expression in oocyte differentiation genes, including *Figa* and *Sohlh2*, and higher expression in genes regulating apoptosis and cell death. These profiling results suggest that cluster 13 may represent a transient stage in oocyte differentiation that later undergo cell death. This hypothesis is supported by the higher level of expression in the genes involving ECM and cytoskeleton organization in cluster 13 germ cells that reflects their activity in transporting organelle and cytoplasmic content. Thus, oocytes may be selected in clusters 4 and 5 germ cells that show difference primarily in cell cycle-related gene expression.

The molecular profiling results of female germ cells are consistent with our proposed model that oocyte determination may be a gradual process that is caused by cytoplasmic enrichment, instead of being pre-determined by a certain molecular profile. This process may involve an initial cytoplasmic transport between two connected germ cells, designating the germ cell that donates cytoplasm to be eliminated. The germ cell with enriched cytoplasm may further communicate with the remaining connected germ cells to gradually transport cytoplasm into branching germ cells. Our results are consistent with recent single-cell RNA-seq data (68), showing that nurse cells detected by lysotracker are gradually activated during oocyte differentiation. The nurse cells and cluster 13 germ cells identified in our present study both have low levels of *Tex14* expression and, based on GO analysis, are enriched in biological processes involved in apoptosis, programmed cell death, regulation of cellular component organization, regulation of organelle organization, and movement of a cell or subcellular component (Supplementary Table 13-15).

In summary, our present study identifies branched germline cyst structures during mouse gametogenesis and differential patterns of cyst fragmentation between female and male cysts in fetal gonads. The patterns of cyst fragmentation, germ cell loss, and distribution of organelle/cytoplasmic content in female cysts indicate that branching germ cells differentiate into primary oocytes within the germline cysts. E-cadherin is involved in branched cyst formation, which may influence primary oocyte number and organelle content in the primary oocytes. Our study sheds light on how the size of the ovarian reserve is determined during mammalian ovary formation.

## Materials and Methods

### Animals

CAG-creER (004682) (Hayashi and McMahon, 2002), R26R-YFP (006148)(Srinivas et al., 2001) and Cdh1/E-cadherin-flox (005319) (Boussadia et al., 2002) mouse strains were acquired from the Jackson Laboratory. Figα-cre mice were obtained from Dr. Jurrien Dean’s lab at the National Institute of of Diabetes and Digestive and Kidney Diseases. All mice were maintained at C57BL/6 background and housed and bred according to the protocol approved by the Institutional Animal Care & Use Committee (IACUC) at the University of Michigan (PRO00008693), the Buck Institute for Research on Aging (A10207) and the University of Missouri (36647). Mice euthanasia in this study was performed by strictly following the protocol from the IACUC. The carbon dioxide (CO_2_) inhalation system in the vivarium facility was used for euthanasia. During euthanasia, the mice were placed into a chamber, in which CO_2_ was added slowly to increase its concentration. The personnel performing euthanasia was required to monitor the process, and wait for at least one minute after no movement, visible inhaling and heartbeat was detected. An approved secondary method was used to ensure the death of the mice.

### Single primordial germ cell lineage labeling and E-cadherin knockout

Single-cell lineage tracing was carried out by using the protocol published in our previous study (Lei and Spradling, 2013, Lei and Spradling, 2016, Lei and Spradling, 2017). Briefly, a single dose (0.4 mg per 40g body weight) of tamoxifen was administered to female (R26R-YFP/YFP) mice that were plugged by male (CAG-creER+/-; R26R-YFP/YFP) mice at E10.5 by intraperitoneal injection. Fetal ovaries and testes were collected at E12.5, E14.5, E17.5 and P0 and lineage-labeled clones were revealed by YFP antibody staining. For single SSC lineage labeling in the adult testis, adult (4 weeks and older) male Pax7-creER+/-; R26R-YFP/YFP mice were injected with a single dose of 5.0 mg tamoxifen by intraperitoneal injection to sparsely label Pax7+spermatogonia stem cells. Testes were collected at day 1, 10, 15 or 20 days after the injection and stained with a YFP antibody for analyzing clone size and cyst structures. To knockout E-cadherin in the single germ cell clone, a single dose (0.8 mg per 40g body weight) of tamoxifen was administered to female (R26R-YFP/YFP; E-cadherin-flox/flox) mice that were plugged by male (CAG-creER+/-; R26R-YFP/YFP; E-cadherin-flox/flox) mice at E10.5 by intraperitoneal injection. Germ cell clones and cysts with Cre recombinase activity and E-cadherin depletion were identified by YFP expression in the germ cells.

### Whole-mount immunostaining

Whole-mount immunostaining for fetal gonads and seminiferous tubule in adult testis were performed as described previously (Lei and Spradling, 2013, Nakagawa et al., 2010). Fetal gonads and adult testes were dissected in PBS and then seminiferous tubules were untangled. Tissues were fixed immediately in cold 4% paraformaldehyde for 1hr (E12.5 and E14.5 gonads and seminiferous tubules) or 2hr (E17.5 and P0 gonads). After washing in PBST_2_ (PBS with 0.1% Tween-20 and 0.5% Triton X-100) for 1 hour, the tissues were incubated at 4°C overnight with primary antibody in PBST_2_ with 10 % donkey serum, 10% BSA and 100 mM Glycine. Tissues were then washed three times with PBST_2_ for at least 15 minutes each, followed by the incubation with the secondary antibodies in PBST_2_ overnight. After washed with PBST_2_, tissues were stained with DAPI in PBS to visualize nuclei. The stained tissues were placed on the coated glass slides (VWR, #48311-703) and mounted on slides with mounting medium (Vector Labs). The stained tissues were imaged using confocal microscopy (Zeiss LSM7000) and analyzed using Image-J and Imaris software (Bitplane). The following primary antibodies and the dilutions (in parentheses) were used in this study: anti-GFP chicken polyclonal antibody (Aves Labs, GFP-1020, 1:1000), anti-TEX14 rabbit polyclonal antibody (ProteinTech, 18351-1-AP, 1:200), anti-RacGAP1 monoclonal antibody (Santa Cruz biotechnology, sc-271110, 1:200), anti-GM130 (BD biosciences, 610822, 1:200), anti-gamma tubulin (abcam, ab179503, 1:200), anti-E-cadherin rat monoclonal antibody (Takara, M108, 1:500), anti-GFRα1 goat polyclonal antibody (R&D Systems, AF560, 1:200), anti-PLZF mouse monoclonal antibody (EMD Millipore, OP128, 1:100), anti-UTF1 rabbit polyclonal antibody (abcam, ab24273, 1:500), anti-KIT rat monoclonal antibody (BD Biosciences, 553352, 1:200), anti-DDX4 rabbit polyclonal antibody (abcam, ab13840, 1:400) and anti-SCP3 rabbit polyclonal antibody (abcam, ab15092, 1:200). The following secondary antibodies from Invitrogen and Jackson ImmunoResearch Inc. were used at the dilution of 1:500: Alexa 488-conjugated donkey anti-chicken (703-546-155), anti-rabbit (715-546-152), anti-rat (712-546-153), anti-mouse (#15-546-151) IgGs, Alexa Rhodamin-conjugated donkey anti-rabbit (711-295-152), anti-rat (712-295-150), anti-mouse (715-296-151) IgGs; and Alexa 647-conjugated donkey anti-rabbit (711-606-152), ant-rat (712-605-153), anti-mouse (715-605-151) IgGs.

### Germline cyst structure analysis

Germ cell clones derived from individual PGCs were imaged at 0.8-1.0 μm per scan. The location of the bridges and germ cells were layout manually by following the location of YFP+ germ cells and RacGAP+ or TEX14+ intercellular bridges using the Image-J image process software. To analyze the number of bridges per spermatocyte in the adult testis, bridges attached to each labeled cell expressing SCP3 are counted.

### Quantification of germ cell volume and Golgi complexes in the germline cyst

Lineage-labeled fetal ovaries stained with GFP, GM130 and TEX14 antibodies, lineage-labeled clones were imaged using confocal microscopy at the Z=1 μm per scan. To measure germ cell diameter of the germ cells in each clone. Two vertical diameters of the biggest cross section of YFP+ germ cells were measured using Image J, and the average value (Rave gc) was used to represent the diameter of the germ cell. The volume of the germ cells (Vgc) was calculated using the equation: Vgc=4/3*3.14*Rave gc^3. To measure the volume of the Golgi complex in each germ cell, ovaries were collected at E17.5 and E18.5 for cysts at the late stage of organelle transport. The volume of the Golgi complex in each germ cell was measured using the surface tool of Imaris software. To compare the volume of Golgi complexes among the germ cells of the cyst, the volume of the Golgi complex in each germ cell was normalized by dividing its Golgi volume by the smallest volume of the Golgi complex in the cyst.

### Measurement of the angles between intercellular bridges

To measure the angles between two adjacent intercellular bridges in the 4-cell cysts, lineage labeled ovaries were collected at E12.5 and stained with GFP and RacGAP antibodies. The labeled germ cell clones were imaged with a LSM 700 confocal microscope using a 40× objective. The angel of the triangle comprised of three points (two adjacent intercellular bridges, and the center of the germ cell nucleus) was measured on Z-stacks of cells using Imaris software.

### Follicle quantification, oocyte volume measurement and B-body quantification

Postnatal day 4 ovaries were collected from the wildtype (Figa-cre+/-; E-cad+/+) and homozygous mutant (Figa-cre+/-;E-cad flox/flox) mice. The ovaries were fixed in cold 4% paraformaldehyde for 2 hrs at 4°C. After fixation, ovaries were washed in PBS and incubated in 30% sucrose overnight before embedding in optimal cutting temperature (OCT) compound. Serial sections were cut at 10 μm of the entire ovary. Sections were stained by using VASA antibody to reveal oocytes. Follicles were counted on every fifth section. The number of follicles per ovary were calculated as follicles/section × total sections. To measure primary oocyte volume and the percentage of the oocytes containing a B-body, wildtype and homozygous P4 ovaries were collected for wholemount antibody staining. The ovaries were stained with VASA, GM130 and E-cadherin antibodies and imaged using confocal microscopy at the Z=1 μm per scan. An area of 100 μm wide and 100 μm deep to the ovarian surface in the ovary was chosen for analysis, and two optical sections with at least 15 μm interval from each ovary were analyzed. Primary oocytes were recognized by VASA positive oocytes that are surrounded by a single layer of flatten pregranulosa cells. For wildtype ovaries, all primary oocytes in the area were analyzed. For homozygous mutant ovaries, only the primary oocytes with E-cadherin negative staining were analyzed. To measure primary oocyte diameter, two vertical diameters of the biggest cross section of each germ cells were measured using Image J, and the average value (Rave gc) was used to represent the diameter of the germ cell. The volume of the germ cells (Vgc) was calculated using the equation: Vgc=4/3*3.14*Rave gc^3. The B-body is defined as a circular Golgi complex in VASA positive primary oocytes. To quantify the percentage of primary oocytes containing a B-body, the number of total oocytes in the above defined area in the wildtype ovary and the number of the oocytes containing a B-body were quantified. For homozygous mutant ovaries, E-cadherin negative oocytes and the ones containing a B-body in the above defined area were quantified.

### Mathematical modeling of the loss of germ cells and intercellular bridges in germ cell clones

The model was written in MATLAB using undirected graph algorithms in which each cell is considered a “node” and each connection or bridge is considered an “edge.” Adjacency matrices describing the connectivity of the cells in cysts were created using Cytoscape for 41 E14.5 clones; all remaining adjacency matrices were created manually. Simulations of two models describing the mechanisms of cyst fragmentation were performed: random cell death and random bridge breakage. Using each clone from E14.5 as an initial model input sample, 250 trials were run; this number was optimized by allowing the ratio of variance to mean for each of the measured statistics to approach a constant. Each mechanism was tested to compare the resulting population of hundreds of simulated broken-down cysts to the experimental E17.5 cyst structure data. There were 41 E14.5 and 29 E17.5 experimental input samples available for the model. The approach to test different fragmentation mechanism of E17.5 compared 29 E17.5 experimental input clones to 22 P0 clones. Many summary statistics of the original (*In Vivo*) and simulated (*In Silico*) cyst networks help to quantitatively measure the size of the cell populations. The connectivity of the cells is given by the degree distribution, where the degree of a cell is the number of bridges it has (or the number of other cells to which it is connected) and the degree distribution is the percentage of cells in a clone with zero, one, two, three or four bridges. Then the averages of each cell number are calculated by multiplying with total cell numbers collected in *In Vivo*. For each summary statistic, a two-sample t-test with 0.05-significance level was used to test the null hypothesis that the data came from independent random samples from normal distributions with equal means and equal but unknown variances.

### Single-cell RNA sequencing

Ovaries were collected at E14.5 and P0 in cold PBS. The tissues were then processed using Neural Tissue Dissociation Kits (MACS Miltenyi Biotec 130-092-628) to dissociate into single cell suspension. The cell suspension was filtered through 40 μm cell strainer to remove the debris. The cells were processed by single-cell 3’ RNA sequencing using the Chromium Single-Cell Controller (10x Genomics, CA) and sequenced by the Advanced Genomics Core of the University of Michigan. The raw sequencing data were processed using 10X Genomics Cell Ranger pipeline and the resulted filtered_feature_bc_matrix of each sample was read into R v3.6.3 and analyzed using Seurat package v3.1.5.

### Statistics

All data was presented as mean ± SD. Nonparametric t tests were run to analyze the difference between two experimental groups. Multiple experimental groups were analyzed using one-way ANOVA. P value level of at least P<0.05 was considered to be statistically significant.

## Acknowledgments

Research reported in this paper was supported by the National Institute of General Medical Sciences (R01GM126028). Dr. Ikami was supported by Toyobo Biotechnology Foundation (Grant in-aid for academic year), Uehara Memorial Foundation Research Fellowship, and the Overseas Research Fellow Award by the Japan Society for the Promotion of Science. We thank Dr. Guy Riddihough at Life Science Editors for editing the manuscript.

